# Hybrid Amyloid Quantum Dot Nanoassemblies to Probe Neuroinflammatory Damage

**DOI:** 10.1101/2023.08.30.555592

**Authors:** Wesley Chiang, Jennifer M. Urban, Francine Yanchik-Slade, Angela Stout, Bradley L. Nilsson, Harris A. Gelbard, Todd D. Krauss

**Affiliations:** Department of Chemistry, Rochester, New York 14627-0216, United States; The Institute of Optics, Rochester, New York 14627-0216, United States; Department of Biochemistry and Biophysics, University of Rochester Medical Center, Rochester, NY, 14642; Center for Neurotherapeutics Discovery and Department of Neurology, University of Rochester Medical Center, Rochester, NY, 14642; Departments of Pediatrics, Neuroscience, and Microbiology and Immunology, University of Rochester Medical Center, Rochester, NY, 14642

**Keywords:** quantum dots, neuronal imaging, biomimetic, neurotoxic oligomers, fluorescence microscopy, amyloid, Alzheimer’s

## Abstract

Various oligomeric species of amyloid-beta have been proposed to play different immunogenic roles in the cellular pathology of Alzheimer’s Disease. However, investigating the role of a homogenous single oligomeric species has been difficult due to highly dynamic oligomerization and fibril formation kinetics that convert between many species. Here we report the design and construction of a quantum dot mimetic for larger spherical oligomeric amyloid species as an “endogenously” fluorescent proxy for this cytotoxic species to investigate its role in inducing inflammatory and stress response states in neuronal and glial cell types.

---

Despite decades of research into the underlying mechanisms that give rise to Alzheimer’s Disease (AD), no clear consensus has emerged as to which cellular phenomenon – amyloidosis, tauopathy, inflammation, oxidative stress – truly drives cognitive impairment.^2^ Based on neuropathologic and genetic evidence,^3^ deposition of amyloid-b (Ab) peptide in the brain gave rise to the amyloid cascade hypothesis in which various oligomeric species of Aβ, with a particular focus on spherical aggregates,^4-6^ have been implicated as being drivers of neurotoxicity.^7-10^ This has heavily influenced development of therapeutic interventions, many of which have targeted metabolism and antibody-mediated removal, with antibodies demonstrating varying degrees of efficacy in Ab plaque removal.^3^

Unfortunately, many of these therapies have demonstrated limited clinical efficacy, despite measurable declines in overall amyloid burden. The disheartening results of these may be explained by the observation of Ab deposition decades before any signs of cognitive decline become clinically apparent.^11^ However, the lack of a strong correlate between decreasing amyloid burden and reduction of cognitive impairment does not imply that simply targeting other molecular drivers, such as tau, will be an effective alternative route for therapy. Indeed, even tau-targeted immunotherapies have had limited clinical efficacy thus far.^12, 13^ The limited success of either therapeutic approaches in conjunction with the complexity of cellular milieus in both neurons and glia in AD pathophysiology may imply that it is a sequence of molecular events that drive cognitive impairment, rather than a sole malefactor.^2, 14^ For example, the correlation of both amyloid and tau accumulation to cognitive decline may tbe linked via dysregulation of both pro- and anti-inflammatory signaling leading to acute neuroinflammatory hallmarks that best correlate with cognitive impairment.^15-19^ In line with such a hypothesis, longitudinal positron emission tomography (PET) studies with Ab and tau stable radioisotopes demonstrated that tau burden correlated with cognitive decline while Ab burden correlated only with tau burden and not cognitive decline.^20^). Taken together, the evidence in aggregate suggests that Ab may initiate tau pathology, and the associated burden of each drives inflammation at different stages within AD progression to mediate cognitive decline.

Thus, understanding the mechanisms by which Ab can initiate progressive tauopathies and associated inflammatory dysregulation will aid in the development of future therapies. It is for these reasons that we have used the following design of a quantum dot biomimetic for spheroidal Aβ oligomers (ABOs), the most commonly implicated neurotoxic species of Aβ,^4-6^ as a tool to interrogate multiple pathways for aberrant neuroimmune signaling and by extension, in future studies, using a similarly constructed tau biomimetic to investigate spreading tauopathies. These tools have the advantage of allowing us to study in situ signaling at the cellular and subcellular level, with the potential to study nanoscale events that may reveal further clues as to how Ab can initiate spreading tauopathies.

However, a well-known potential disadvantage of using QDs as biological imaging probes is that they often have drastically different physical characteristics (i.e. size, valency, target density, etc.) compared to the biological target molecule they are labeling.^21, 22^ Indeed, functionalization of QDs with biomolecules generally results in the formation of large, spherical probes presenting many copies of the biomolecule of interest on the QD surface. Based on differences in size and ligand valency, it is very reasonable to assume that QDs may behave differently than the ligand alone in a biological context. Thus, when using QD probes it is common to focus on overall cytotoxicity or biodistribution of QDs *in vitro* and *in vivo*.^23-29^ However, it is far less common to investigate whether the biological system behaves appropriately in response to the biomolecular probe that is tethered to the surface of the QD. For example, the aforementioned concerns of QD-based biological probes may present themselves as advantageous if applied as a biological mimetic, rather than a simple biological label. Specifically, the size and valency of ligand-functionalized QDs can be tailored to mimic endogenous spherical biological macromolecules, this can range from spherical oligomeric proteins to virus-like nanoparticles to even lipid vesicles.^30^ As such, we posit that coating the QD surface with features of organic matter resembling aggregated macro-molecules, such as AβOs, a QD-based biomimetic may be constructed to be structurally and functionally perceived by cells akin to endogenous spheroidal AβOs (Figure 1). These spheroidal ?βO-mimicking QDs (ABQDs) may serve as nanoscale biophysical tools to determine the mechanistic role of spherical AβOs in mediating neuroinflammation and subsequent tauopathies in AD pathology.

**Figure 1.**
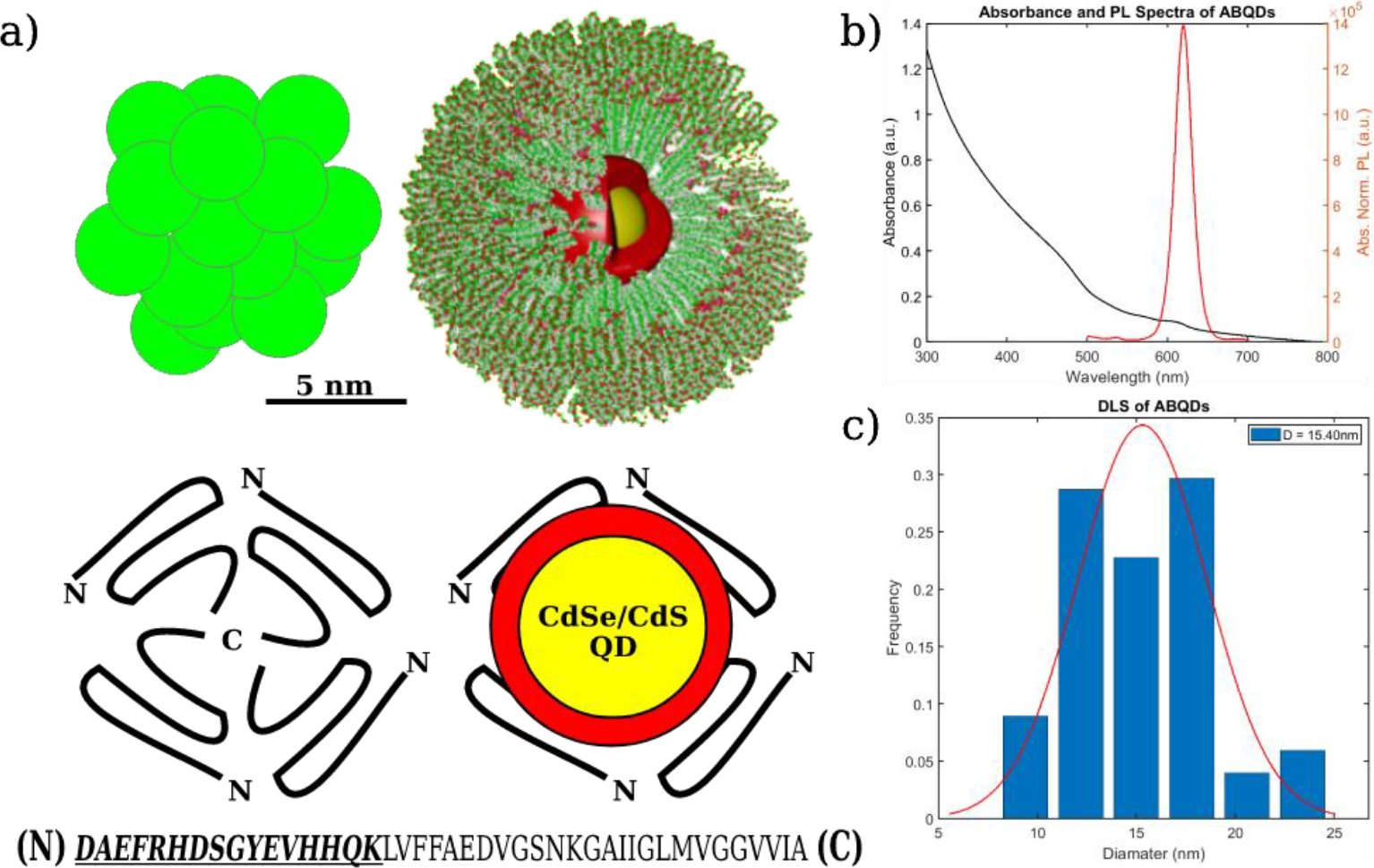
Diagram of conceptual framework behind ABQD design and characterization of optical and size properties. **(a)** Size comparison of spherical aggregates of amyloid-beta 42 peptides to a CdSe/CdS QD encapsulated in a DSPE-PEG2k micelle (top), proposed oligomerization of peptides by Ahmed *et al*. compared to structural design of ABQDs (middle), where the first 16 amino acids of the N-terminus are synthesized (below) and used to functionalize the micellar surface.^1^ **(b)** Confirmation that ABQDs retain the optical properties of the original CdSe/CdS QDs. **(c)** DLS characterization of ABQD size distribution, with average hydrodynamic radius ≈15.40*nm*.

Herein, we report the construction of ABQDs via micelle encapsulation of CdS QDs in polymerized phospholipids decorated with 16-residue peptide sequences of Aβ(1-16). Functional mimicry of the ABQD constructs was assessed by measuring inflammatory hallmarks and calcium signaling in primary cultures of hippocampal neuroglia isolated from Sprague-Dawley rats. We also observed changes in phagocytic activity and activation of a pro-inflammatory and endoplasmic reticulum (ER) stress responses in an immortalized microglial (BV-2) cell line. Finally, we show that these effects are attenuated by co-treatment with a broad-spectrum mixed-lineage kinase inhibitor, URMC-099, previously demonstrated to ameliorate these pathologic hallmarks with *in vitro* and *in vivo* models of AD.^31, 32^ In aggregate, our results highlight the development of a new, robust fluorescent tool to mimic spherical aggregates of Aβ42 that can be utilized to elucidate neurotoxic mechanisms related to AD and the effectiveness of therapeutic interventions in mitigating these effects.

## RESULTS AND DISCUSSION

### ABQDs structurally resemble spheroidal AβOs

Our proposed construct models the assembly of Aβ42 oligomers such that the N-terminal amino acid residues are exposed to the biological environment, serving as the antigenic region responsible for biomolecular recognition and signal transduction.^4-6^ Thus, we synthesized, via solid-phase peptide synthesis, the first 16 amino acids from the N-terminus of Aβ42 and attached the peptides to a phospholipid-PEG construct (DSPE-PEG_2k_) via a cysteine-maleimide conjugation scheme (Scheme S1). The successful synthesis of the peptide and conjugation to form an Aβ(1-16) functionalized lipid PEG (DSPE-PEG_2k_-Aβ) was characterized via analytical high performance liquid chromatography (HPLC) and matrix-assisted laser desorption ionization time-of-flight (MALDI-ToF) mass spectrometry (Figure S1).

However, beyond ensuring that the approach to decorate the QD surface appropriately modeled the exposed region of spheroidal AβOs, detailed characterization of the structural resemblance of ABQDs to endogenous A β42 spheroidal oligomers, as diagrammed in Figure 1a, is key to ensuring that the ABQDs can be used as a functional proxy. Thus, after micellar encapsulation of CdSe/CdS with DSPE-PEG_2k_-Aβ to form ABQDs, the heterogeneous population of mimetic constructs were separated into unique size fractions. Each size fraction was measured with dynamic light scattering (DLS) to identify the fraction that best fit the reported diameter of neurotoxic spheroidal amyloid oligomers, previously reported to be approximately 12 nm -20 nm.^1, 4, 5, 8^ We characterized absorbance and photoluminescence parameters to ensure that desirable optical properties of QDs are retained (Figures 1b, c). The absorbance spectra (black line) in Figure 1b exhibit low signal past the lowest energy transition of the QD (> 620nm) in Figure 1b that would arise from scattering from empty micelles. Complemented by the DLS data, in Figure 1c, showing a size distribution of 10 nm -22.5 nm, the isolated size fraction of interest contains CdSe/CdS encapsulated Aβ(1-16) functionalized micelles that structurally resemble spheroidal AβOs. Specifically, the endogenous structures have been reported to range from 11 nm – 17.5 nm and purportedly assemble where the C-terminal domains of monomers interact into structured globular domains while the 16 N-terminal amino acids are flexible and exposed on the outer region of these aggregates..^1, 5, 6^

### ABQDs recapitulate Aβ42-associated damage phenotypes in neurons and astrocytes

After validating that the ABQDs have been constructed to mimic key structural parameters (size, general shape, and surface amino acid sequence) of endogenous spheroidal AβOs, we exposed primary cultures of rat hippocampal neurons and astrocytes to 50 nM of ABQDs. Using immunocytochemical labeling of neuronal dendrites with microtubule-associated protein 2 (MAP2; magenta) and astrocytes with glial fibrillary acidic protein (GFAP; white), (Figure 2a), we were able characterize the functional capacity of the ABQDs to induce the presence of neuroinflammatory hallmarks, such as dendritic beading (Figures 2b, c) and astrogliosis (Figure 2d). Dendritic beading, defined as the observation of focal swellings in postsynaptic neuronal processes (Figure 2c), is a hallmark of synaptic injury and neurite damage commonly observed in AD neuropathology.^33-37^ Additionally, astrogliosis arises from the activation of a reactive, pro-inflammatory state in astrocytes typified by increased spatial distribution and expression levels of astrocytic cytoskeletal components, such as GFAP.^38-40^ While astrogliosis is a general hallmark of neuroinflammation, direct modulation of astrocyte activation and phenotype has been implicated in toxic AβO function.^40-42^ Comparatively, treating neuroglial cultures with 50 nM of micelle encapsulated quantum dots lacking Aβ(1-16) functionalization (PEGQDs), results in little or no dendritic beading compared to vehicle treated groups (Figure 2b), but a small, yet significant, induction of astrogliosis (Figure 2d). We attribute this discrepancy in the activation of inflammatory profiles in neurons versus astrocytes to arise from the recognition of PEGQDs by astrocytes as non-specific extracellular debris that results in phagocytosis or endocytosis of PEGQDs by astrocytes (Figure S3). This associated astrogliosis may be seen as homeostatic regulation of the neuronal environment.^43-46^ However, astrogliosis is likely exacerbated by an amyloid specific interaction with the ABQDs, potentially mediated through complement dependent recognition of Aβ at synapses.^40, 42, 47-53^ Furthermore, progressive accumulation of Ab and tau in 5X-FAD and JNPL3 mice respectively can induce reductions in PSD-95 in apical hippocampal dendrites, with a similar phenomena in hippocampal AD brain sections.^54^ When examining the association of ABQDs compared to PEGQDs to neurons, we see that without the functionalization of Aβ(1-16) on the micellar surface, PEGQDs are unable to bind to post-synaptic densities (PSD95; green), while ABQDs exhibit specific association to only PSD95 of neurons in the cultures (Figures 2e-g). Therefore, the neuronal damage mediated by the ABQDs is due to an amyloid-dependent biomolecular interaction with the neurons.

**Figure 2.**
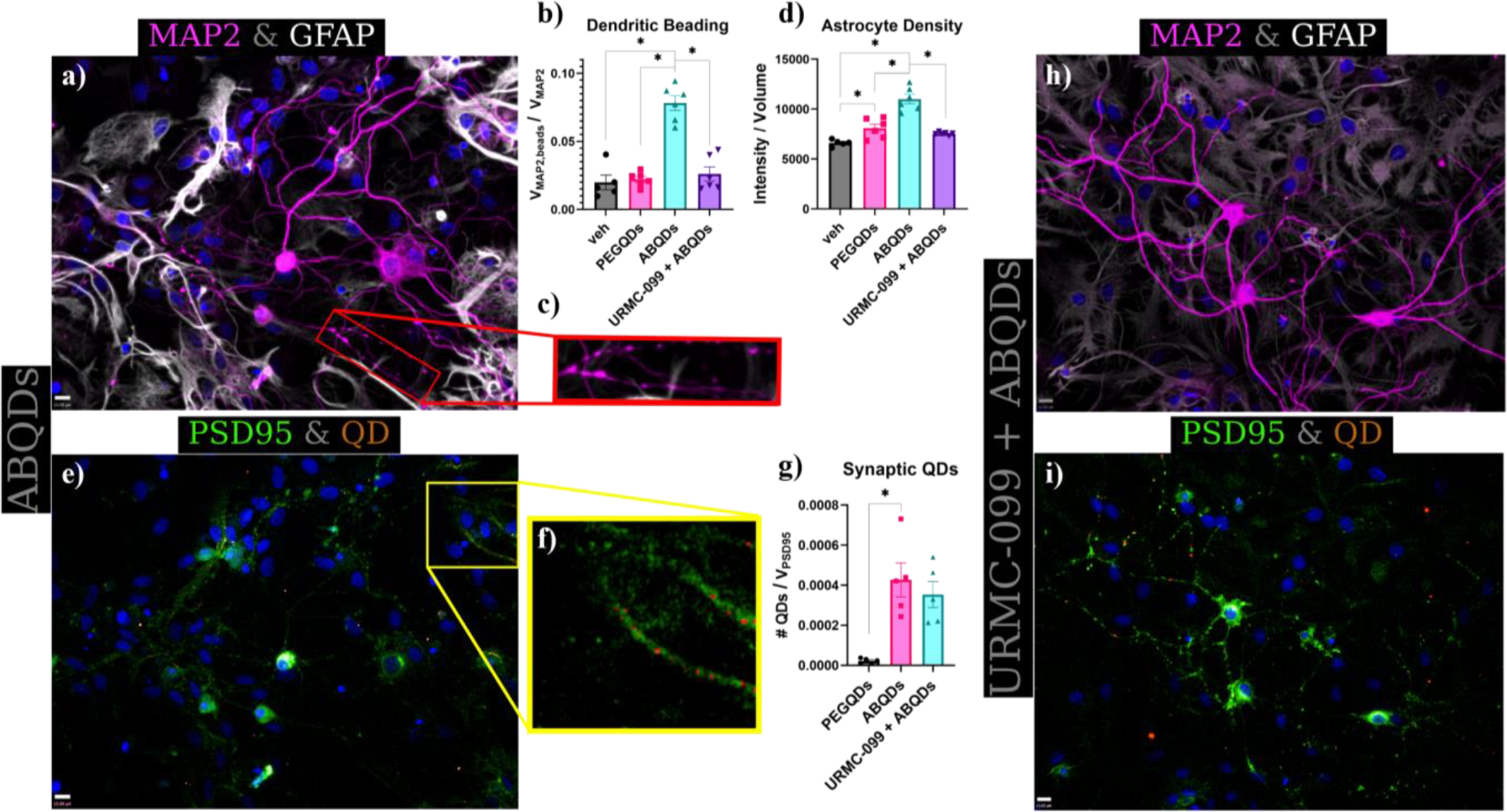
Neuroinflammatory activation by 50 nM of ABQDs that is mitigated by co-treatment with 100 nM of URMC-099. Immunofluorescent labeling of MAP2 and GFAP demonstrate the formation of focal swellings along neuronal processes (i.e. dendritic beading) complemented by astrogliosis in ABQD only treated neuroglial cultures **(a, c)**; co-treatment with URMC-099 reduces the observation of such inflammatory hallmarks **(h)**. In both treatment groups, ABQDs are shown to selectively colocalize with PSD95, indicative of synaptic targeting and recognition commonly associated with amyloid pathology **(e, f, i)**. Statistical analyses of these observations are performed using one-way ANOVA with a Holm-Sidak post-hoc correction (*; p < 0.05) as plotted in **(b, d, g)**.

### URMC-099 attenuates ABQD induced inflammation

In previous work, we have shown that a small-molecule kinase inhibitor against mixed-lineage kinases (MLKs), known as URMC-099, acted in an anti-inflammatory and neuroprotective manner in murine models of various neuroinflammatory and neurodegenerative disorders.^31, 55-59^ While most of these studies have been conducted in either murine models, or in cultures focused more on microglia, URMC-099 may also elicit protective effects directly in neurons and astrocytes, given the ubiquitous expression and subsequent role of MLKs in nearly all eukaryotic cell types.^60^ As such, we co-treated the neuroglia cultures with both 50 nM ABQDs and 100 nM URMC-099 to see if a direct inhibition of pro-inflammatory kinase signaling in neurons and astrocytes would reduce the manifestation of neuroinflammatory hallmarks. In agreement with our hypothesis of direct inhibition of MLKs, we saw a reduction in the amount of dendritic beading and astrogliosis in the co-treated cultures (Figures 2b, d, h). This neuroprotection was not due to URMC-099 mediated disruption of ABQD recognition of synaptic targets, given that we observed no significant (one-way ANOVA + Holm-Sidak post-hoc; p > 0.05) difference in the fraction of ABQDs co-localized with PSD-95 in either treatment groups (Figures 2g, i).

### ABQDs induce changes in neuronal calcium transients

Previous studies have linked AβOs to excitotoxicity in neurons due to dysregulation of intracellular calcium transients, leading to the neuroinflammatory hallmarks of synaptic and dendritic damage that we observed in Figure 2.^33, 61-63^ To further validate the ability of ABQDs to induce an AβO-associated patho-physiological response in neurons, we examined dysregulation of homeostatic calcium transients in response to the ABQD constructs. Neuronal cultures were treated with an equivalent dilution of nanopure water (vehicle), 10 nM PEGQDs, or 10 nM of ABQDs for 10 minutes, followed by induction of a calcium transient after treatment with 30 mM KCl. As shown in Figure 3, the vehicle and PEGQD treated cultures have no observable difference in the magnitude of calcium transients, as measured by the total fluorescent intensity normalized to the area of interest. In contrast, ABQDs induce a much stronger calcium response (1.5x to PEGQDs and 1.75x to vehicle), observable at the 200 sec frame (Figure 3c) and the normalized intensity shown in Figure 3d. This increased calcium influx in the ABQD-treated neuronal cultures suggests excitotoxic stress in neurons with observed neurite damage.

**Figure 3.**
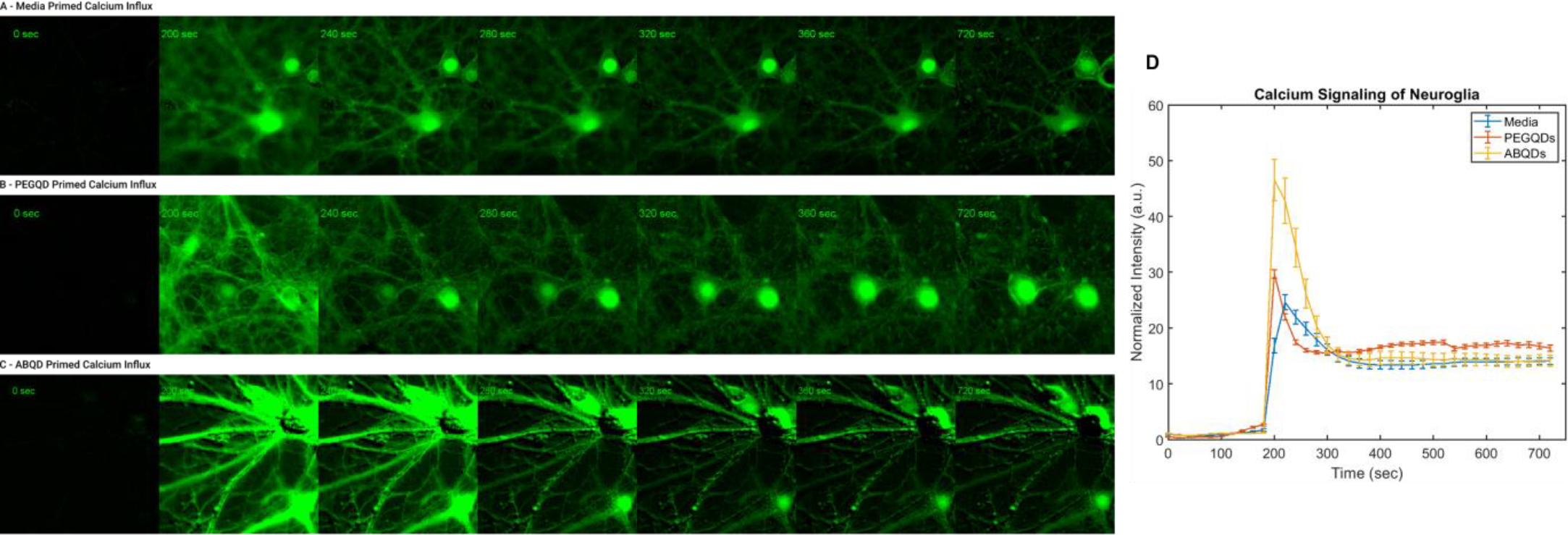
Calcium transients reflecting amyloid-dependent response from neuroglial cultures. Co-cultures of neurons and astrocytes were primed with just an equal volume vehicle **(a)**, 10 nM of PEGQDs **(b)**, or 10 nM of ABQDs **(c)** followed by 30 mM KCl calcium induction. The results of the induced calcium transients are plotted in **(d)** and reflect the enhance calcium spike from ABQD treated cultures compare to media and PEGQD treatments.

### Microglia exhibit amyloid-specific phagocytic response to ABQDs

Beyond the direct neurotoxic effects of ABQDs on neurons and astrocytes, we characterized the capacity for ABQDs to interact with microglia in an amyloid-dependent manner that occurs in murine models of AD. Specifically, microglia play an essential role in phagocytic uptake and clearing of extracellular amyloid to ameliorate AβO-associated pathology. However, when the amyloid burden is large, microglia processing of AβOs leads to activation of an ER stress state associated with an unfolded protein response (UPR) and subsequent pro-inflammatory response.^31, 64-66^ Thus, we treated an immortalized microglial cell line (BV-2) with either 50 nM PEGQDs (Figure S4) or 50 nM ABQDs (Figure 4) and examined the differential uptake and intracellular processing of these two constructs.

**Figure 4.**
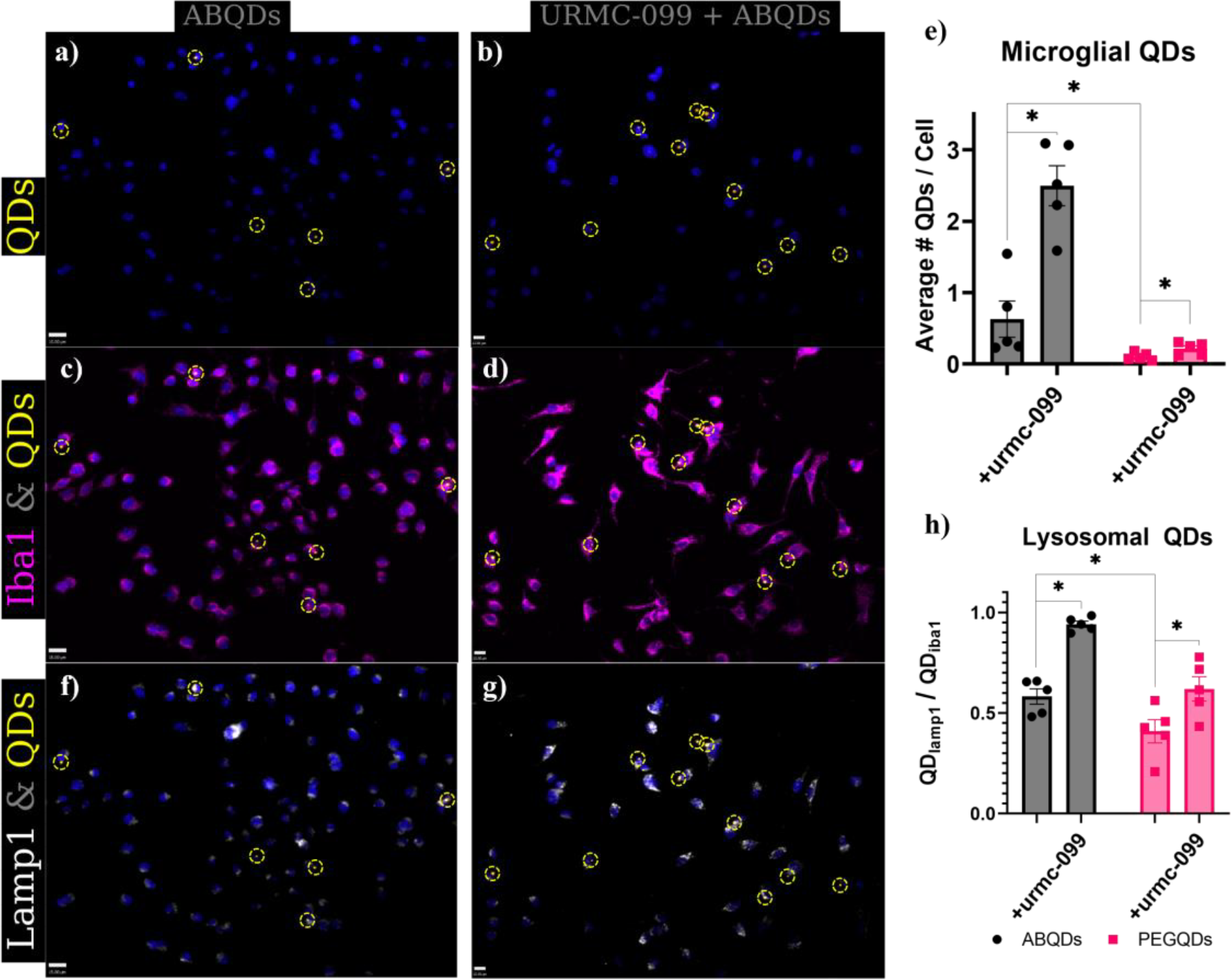
Amyloid-dependent uptake and autophagy of ABQDs. Co-treatment of BV-2 cultures with 100 nM URMC-099 increased the observed ABQD-puncta associated with Iba1+ cells compared to cultures treated with 50 nM ABQDs alone **(a-d)**. This increase of phagocytosis by microglia was observed in both ABQD and PEGQD treated groups (two-way ANOVA + Holm-Sidak post-hoc; * p < 0.05), but the raw numbers of microglia associated in 50 nM PEGQD were considerably negligible. The increase in phagocytosis by URMC-099 co-treatment also resulted in creased autophagy, noted by increased colocalization with Lamp1+ regions in the BV-2 **(f-h)**.

Without the presence of Aβ(1-16) peptides on the surface, the PEGQDs are observed at much lower frequency, defined by the total number of detected QD puncta normalized to the total number of cells (unique DAPI objects), than the ABQDs are in the BV-2 cultures (Figures S3a and 4a), though in both cases all QD PL is observed to be co-localized with Iba-1 labeled microglia. Specifically, in Figures S3b and 4b, we rarely identified neither ABQD nor PEGQD associated fluorescence that was not coincident with Iba-1 associated fluorescence; these are represented by orange puncta outlined with dashed yellow circles and magenta objects, respectively, in the fluorescence micrographs. Like astrocytes, the differential association of PEGQDs and ABQDs to the microglia likely arises from distinct glial phagocytic and processing pathways for extracellular debris and amyloid. ^10, 31, 44, 46, 64, 66-69^ To test this hypothesis, we treated microglial cultures with 100 nM of URMC-099, which has been previously shown to increase amyloid uptake in microglia, but no substantial increase of polymeric nanoparticles in monocyte-derived macrophages.^31, 64, 68^ Correspondingly, Figures 4b and 4d demonstrate an increased number of observed puncta associated with QD PL that is highly correlated with Iba-1+ microglia. Normalizing the count of QD PL puncta by total number of microglia observed (Figure 4e), there is a statistically significant (p < 0.05, two-way ANOVA + Holm-Sidak post-hoc) increase in ABQD uptake after URMC-099 treatment. While statistical evaluation finds a similarly significant increase in PEGQD phagocytosis, which aligns with the increased autophagic capacity of URMC-099 treated microglia, the raw numbers are essentially negligible in difference; the increase in PEGQDs phagocytosis goes from less than 1 PEGQD phagocytosed per 20 microglia to 1 in 10 microglia, while ABQD phagocytosis starkly increases from ~1 per microglia to ~2.5 per microglia Based on these observations, we conclude that the ABQDs are likely recognized in an amyloid-dependent mechanism that leads to differential phagocytic uptake compared to surface-bare PEGQDs. Importantly, while glial phagocytosis of monomeric amyloid and some fibrillar forms of Aβ42 has been previously reported, phagocytosis of the larger, cytotoxic spheroidal AβOs (such as this ABQD biomimetic) has not been clearly documented, to our knowledge, until this study.^31, 64, 66, 70^

### URMC-099 primes microglial autophagy response to alleviate ABQD-associated pro-inflammatory activation and ER stress

To further assess microglial recognition of ABQDs in an amyloid-specific process, we examined the intracellular fate of ABQDs compared to PEGQDs after phagocytic uptake. Amyloid-dependent activation of microglia should result in a pro-inflammatory state and subsequent increased phagocytic activity, as well as induction of ER stress via an unfolded protein response (UPR).^32, 69-77^ In line with UPR-mediated stress and inflammatory activation, phagocytosis of ABQDs should be ultimately trafficked to either a proteasomal degradation pathway or autophagy-associated lysosomal digestion. Correspondingly, as shown in Figures 4f and 4h, there is a significant (p < 0.05, two-way ANOVA + Holm-Sidak post-hoc) increase the number and fraction of ABQDs in Lamp1 immunostained lysosomes, compared to PEGQDs. Additionally, while the PEGQD treated microglia exhibit a non-negligible fraction of lysosomal association (≥ 50% of observed microglia-associated PEGQDs are also colocalized with Lamp1), the raw count of PEGQDs in lysosomes, and microglia in general, is small, as reflected by the low frequency of microglia-associated PEGQDs (Figure 4e). In line with these observations, we expected that URMC-099 treatment likely biased ABQD processing toward the autophagy pathway and led to the marked (p < 0.05, two-way ANOVA + Holm-Sidak post-hoc) increase in association with Lamp1 lysosomes shown in Figures 4g and 4h. Specifically, previous studies of URMC-099 have characterized increased autophagy in monocyte-derived macrophages, due to increased nuclear translocation of transcription factor EB (TFEB) via a JNK/mTORC1 axis, and have linked this mechanisms as being responsible for increased autophagolysosomal processing of Aβ monomers in microglia.^31, 58, 64, 68^

As confirmation that these phenotypic changes in microglial activity in response to ABQDs is driven by proinflammatory activation and ER stress, we isolated RNA from the BV-2 cells and performed reverse transcription quantitative polymerase chain reaction (RT-qPCR) to examine changes in transcriptional activity of C-X-C motif chemokine ligand 10 (CXCL10) and C/EBP homologous protein (CHOP) as well as differential splicing of X-box binding protein 1 (XBP-1). Changes in intracellular CXCL10 can represent inflammatory activation of the microglia, due to a positive feedback pathway with secreted chemokine in these activated phenotypes that can further recruit inflammatory leukocytes.^74-76^ CHOP is a downstream effector of the PERK-eIF2α ER stress transduction pathway and is a transcription factor associated with apoptosis in many neurodegenerative disorders, such as AD.^71, 72, 78-80^ In a separate, IRE1-dependent, ER stress pathway, IRE1 activation causes a frame shift in the splicing of XBP1, a transcription factor involved in the regulation of other ER regulatory transcripts and proinflammatory cytokine production; changes in the relative fractions of spliced XBP1 (sXBP1) to unspliced XBP1 (usXBP1) would indicate changes in the activation state of the microglia due to this IRE-1 dependent ER stress response. For example, as a transcription factor sXBP1 regulates various biosynthetic pathways necessary to maintain healthy ER function, but excessive ER stress drives it to exacerbate pro-inflammatory signaling associated in AD and other neuroinflammatory diseases.^71-73, 79, 80^

Thus, changes in CXCL10 transcript levels would give insight into the inflammatory state of the BV-2 cells in various treatment groups, while CHOP transcript levels and changes in sXBP1 to usXBP1 populations would provide insight in ER stress activation, as shown Figure 5. In agreement with our hypothesis, we saw a significant (p < 0.05, one-way ANOVA + Holm-Sidak post-hoc) increase of CXCL10 and CHOP transcripts due to ABQD treatment as well as increased splicing of XBP1, represented by the fraction of sXBP1 to usXBP1. Furthermore, these effects were ameliorated with URMC-099 treatment, though not to basal levels observed in the two control groups treated with either vehicle or URMC-099 only. This may be attributed to tightly regulated levels of sXBP1 activation and inflammatory activation involved in homeostatic maintenance by microglia.^72-74^ Of interest, the ABQDs seem to act more on the IRE1-XBP1 axis to result in a heightened inflammatory state, as shown by the large increases in sXBP1 and CXCL10, while only a modest, but significant, increase in CHOP. In aggregate, these observations suggest that BV-2 microglia recognize and process ABQDs in an amyloid-associated stress response mechanism that is attenuated with URMC-099 treatment. While URMC-099 has been previously studied in various amyloid and AD models, its capacity to specifically ameliorate damage against larger oligomeric species has not been demonstrated before. Additionally, data from these ABQDs provide potential insight into the specific ER stress pathway activated by the larger spheroidal oligomers that they mimic.

**Figure 5.**
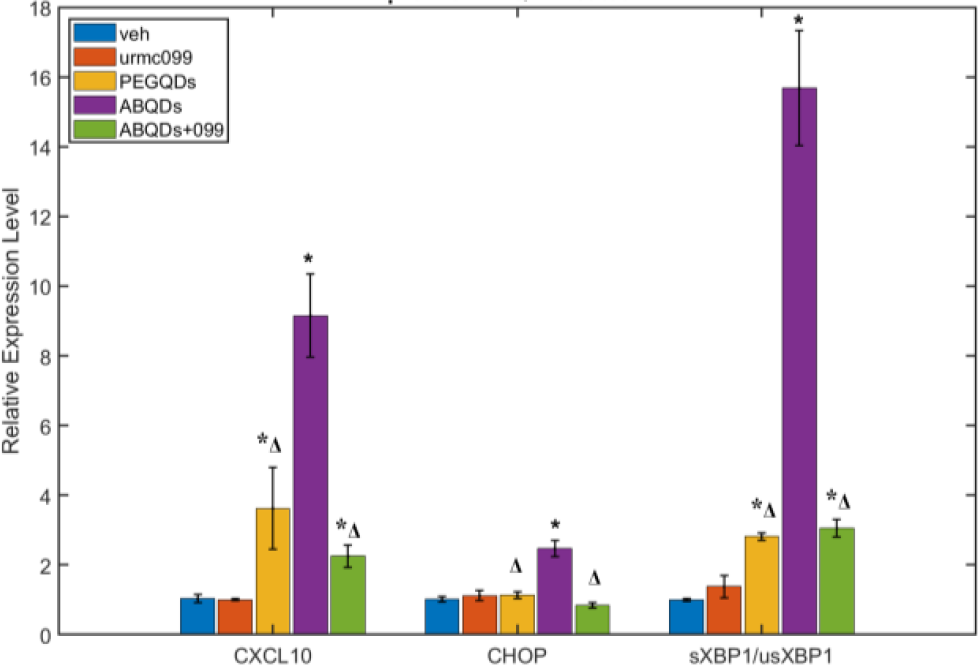
ABQD-mediated ER stress and autophagy signaling in BV-2. CXCL10, CHOP, and splicing variants of XBP1 selected as representative target transcripts for RT-qPCR analysis of inflammatory and ER stress activation specific to ABQDs. * & ^Δ^ denote statistically significant differences (p < 0.05, one-way ANOVA + Holm-Sidak post-hoc) when compared to vehicle or ABQD treatments, respectively.

## CONCLUSION

Understanding the roles of individual oligomeric species of Aβ42 in initiating neuroinflammatory signaling and subsequent pathological events, such as tauopathies, will aid in the precise design of therapeutic strategies to combat AD. In line with this mission, we have designed and constructed a QD-based biomimetic of larger spherical aggregates of AβOs, a proposed cytotoxic initiator of inflammation in AD. The QD core allows this mimetic structure to act as a proxy for an endogenously labeled variant of the native pathologic AβO species, without introducing exogenous labels that may perturb essential aggregation or biomolecular recognition sites. We have validated this ABQD construct to be a structural and functional mimic for the AβOs they mimic and have taken advantage of the enhanced optical properties of the QD core to examine its localization in neurons, astrocytes, and microglia. Lastly, the ABQD constructs are used in tandem with URMC-099 to validate the proper activation of inflammatory and stress signaling response pathways associated with AD. To our knowledge, this work demonstrates for the first time that URMC-099 is able to facilitate increased phagocytosis and autophagy of amyloid species that are extracellularly aggregated. In future work, we hope to apply this tool to further examine the coordination between AβOs and localized neuronal damage, calcium excitotoxicity, and induction of tauopathies at subcellular resolutions.

## EXPERIMENTAL METHODS

For a complete set of materials and methods, please see the associated supplementary document.

### Construction and Characterization of ABQDs

The assembly of ABQDs is comprised of the synthesis of CdSe/CdS QDs followed by encapsulation into a lipid PEG micelle functionalized with Aβ(1-16) peptides. *Synthesis of CdSe/CdS. QDs* were synthesized using a hot-injection protocol adapted from our previous work to produce CdSe cores, followed by growth of CdS shells ^81-83^. The final oleic acid (OA) capped QDs were suspended in a nonpolar solvent, such as toluene or chloroform and spectrally characterized by absorbance and photoluminescence measurements, complemented by transmission electron microscopy. The QDs are stored in glovebox filled with an inert gas, such as N_2_ until needed.

#### Synthesis of DSPE-PEG_2k_-Aβ

The polymerized phospholipid (DSPE-PEG_2k_-Maleimide) used to form the self-assembled micelles were purchased from Avanti Polar Lipids (cat: 880126C-25mg). Before micellar encapsulation, the monomers were conjugated with peptides of Aβ(1-16)-PEG-CG via a cysteine-maleimide reaction. The peptides were synthesized via solid-phase peptide synthesis, purified and quantified via reverse phase HPLC, characterized by MALDI-ToF, and lyophilized for storage until needed. The final construct (DSPE-PEG_2k_-Aβ) was purified via dialysis and characterized using analytical HPLC and MALDI-ToF.

#### Assembly of ABQDs

The CdSe/CdS QDs were encapsulated into micelles formed by self-assembly of DSPE-PEG_2k_-Aβ via a modified dual solvent exchange method. In brief, the QDs and lipid-PEG monomers were both suspended in chloroform and then separately sonicated to minimize the presence of preformed aggregates. The solutions were then combined, vortexed and sonicated, then concentrated using a rotary evaporator. As the solution volume approached that of a gel-like film, the solution was removed from the rotary evaporator, nanopure water was added, and then reconnected to the rotary evaporator. The mixture was kept on the rotary evaporator until all organic phase bubbled off and the QDs were transferred into a single aqueous phase. The final product was purified using a 0.1 μm syringe filter, a size exclusion spin filter, and then a round of ultracentrifugation followed by centrifugation of the resuspended pellet. The final pellet was resuspended and both that and the supernatant were analyzed via DLS to determine which fraction to use.

### Primary neuroglial cultures

Cell cultures for all experiments consisted of primary mixed rat hippocampal neuronal and astroglia cultures at 18-21 days *in vitro* (DIV) grown on either glass or fused silica coverslips.

*Neuroinflammatory Activation* experiments were conducted by exposing the neuroglial cultures to vehicle treatments, 50 nM PEGQDs, 50 nM ABQDs, or 100 nM URMC-099 and 50 nM ABQDs. All treatments were dissolved in neurobasal media (ThermoFisher; cat: 21103049) supplemented with B27 without antioxidants (ThermoFisher; cat: 10889038) and 1% Gluta-Max (ThermoFisher; cat: 35050061). Vehicle treatments were either equivolume dilutions of nanopure water or both nanopure and DMSO in the supplemented neurobasal medium. The cultures were incubated at 37°C and 5% CO_2_ overnight (16-20 hours), before fixation and indirect immunofluorescent staining for imaging. Imaging was conducted in a structured illumination format on an Olympus BX51 microscope equipped with an OptiGrid element. The images were processed with Volocity 3D Image Analysis software (PerkinElmer).

*Calcium Signaling* was performed by priming the neuroglial cultures with a nanopure vehicle dilution, 10 nM PEGQDs, or 10 nM ABQDs all in supplemented neurobasal media. The priming occurred 10 minutes, followed by calcium transient induction with a 30 mM KCl solution in supplemented neurobasal medium. The calcium transients were recorded using an inverted microscope with 100 ms exposure times at 20 sec intervals over 12 minutes. The calcium transients were detected using a fluorochrome, Fluo-4AM (ThermoFisher; cat: F14201). The resultant videos were processed using ImageJ-2 (National Institutes of Health).

**Microglia cultures** were comprised of an immortalized murine microglial cell line (BV-2). Before treatment, the cultures were incubated for 1 hour in a reduced serum condition comprised of Dulbecco’s Modified Eagle Medium (DMEM, ThermoFisher; cat: 10313039) supplemented with 1% fetal bovine serum (FBS, Atlas Biologicals; cat: F-0500-D), 1% GlutaMax, and 1% penicillin-streptomycin (ThermoFisher; cat: 15140122). This was followed by overnight (16-20 hour) incubation at 37°C and 5% CO_2_ of vehicle, 50 nM PEGQDs, 50 nM ABQDs, or 100 nM URMC-099 with either 50 nM PEGQDs or ABQDs all in the same reduced serum medium. The cells were either then fixed and indirectly immunofluorescent labeled for imaging, or RNA was extracted for RT-qPCR analysis.

## ASSOCIATED CONTENT

### Supporting Information

The Supporting Information is available free of charge on the ACS Publications website.

Full experimental procedures, chemical structures, synthetic peptide and quantum dot characterization data (PDF)

Movie S1. Calcium influx due to vehicle treatment (.avi)

Movie S2. Calcium influx due to PEGQD treatment (.avi)

Movie S3. Calcium influx due to ABQD treatment (.avi)

### Notes

HAG is the Chief Science Officer and NT is a member of the scientific advisory board of Pioneura Corp, (Fairport, NY), which holds the exclusive license for URMC-099, but did not contribute either salary support or funding for this work. All remaining authors declare no conflict of interest.

## ACKNOWLEDGMENT

The authors thank Michael Franchot at the UR LLE for creating the artistic rendering of a micelle coated QD in Figure 1, as well as the Integrated Nanosystems Center and the Structural Biology & Biophysics facility at the University of Rochester for providing access to the TEM and DLS instrumentation, respectively. We would also like to thank Madeline Jensen and her mentor Dr. Eric Wagner for their help with troubleshooting the RT-qPCR experiments and access to their qPCR machine. Portions of this work was supported by the National Science Foundation (NSF) CHE-1609365 (TDK) and DMR-1148836 (BLN); National Institutes of Health (NIH) R01HL138538 (BLN), R01MH64570 (HAG), R21NS128502 (HAG and TDK), T32GM135134 (WC), T32GM118283 (FY), and S10OD030302 (MALDI instrumentation); and the University of Rochester Sproull Fellowship (JMU). Additionally, this research was supported by a grant from the University of Rochester Center for AIDS Research (CFAR), an NIH-funded program (P30AI078498). The content is solely the responsibility of the authors and does not necessarily represent the official views of the National Institutes of Health.

