## Supplemental File for "Hybrid Amyloid Quantum Dot Nanoassemblies to Probe Neuroinflammatory Damage"

(INSERT TITLE HERE)

### Materials and Methods

#### CdSe/CdS QD synthesis

*Reagents*: All reagents were used as purchased from the manufacturer without further purification. The following were purchased from Sigma-Alrich: trioctylephosphine (TOP; cat: 718165), selenium pellets (Se; cat: 209643), diphenylphosphine (DPP; cat: 252964), cadmium oxide (CdO; cat: 202894), trioctylphosphine oxide (TOPO; cat: 223301), 1-hexadecylamine (HDA; cat: 445312), oleic acid (OAc; cat: 364525), octadecylamine (ODA; cat: 74750), oleylamine (OAm; cat: O7805), octadecene (ODE; cat: O806), octanethiol (OT; cat: 471836), tetradecylphosphonic acid (TDPA; cat: 736414).

*Precursor Synthesis:* Prior to synthesis of CdSe cores and shelling to generate CdSe/CdS core/shell QDs, both ~1 M TOP:Se and ~0.08 M Cd-oleate were synthesized for each respective reaction. For 1 M TOP:Se, 680 mg Se pellets and 8.5 mL TOP were combined in a scintillation vial and mixed at 60 ^o^C overnight in a glovebox to result in a clear solution. A small amount of secondary phosphines, 90 uL DPP, was added to help promote efficient QD nucleation.^1^ For 0.08 M Cd-oleate, 250 mg CdO, 2.6 mL OAc, and 20 mL ODA were combined in a side-arm storage flask (ChemGlass; cat: AF-0522) and connected to a Schlenk line. The contents were degassed at room temperature, followed by 90 minutes of heating at 270 ^o^C under an inert gas (N_2_) environment. The flask was then cooled to room temperature, with 1.3 mL of OAm injected at 150 ^o^C to help prevent solidification of the product, which was stored in a glovebox until needed.

*CdSe core synthesis:* The CdSe cores were synthesized from an adaptation of procedures previously reported.^2, 3^ To a 3-neck flask was added 820 mg CdO, 16.2 g TOPO, 37 g HDA, and 3.2 g TDPA. The flask was connected to a Schlenk line and purged with N_2_ and then kept under this inert gas environment while the contents were heated to 90 ^o^C. The contents were then degassed by 3 cycles of evacuation (<100 mT) and refiling with N_2_, followed by heating to 320^o^C with rapid stirring. Cd-TDPA formation was visually determined via changes in the transparency of the solution from an opaque to translucent solution. The contents were then cooled to 260 ^o^C, and 8.0 mL 1 M TOP:Se was rapidly injected at this temperature while maintaining the same stir rate. The CdSe cores were grown for 2 to 3 hours until the desired size was achieved, as determined by PL measurements of aliquots taken routinely after the 2 hour timepoint. The reaction was then quenched by removing the flask from heat and applying forced air to bring the solution to 200 ^o^C before submerging the flask into a water bath to rapidly cool the solution to 100 ^o^C. The solution was then injected with 40 mL of ButOH and allowed to cool for ~1 hour before performing several rounds of washing followed by two cycles of size-selective precipitation. The final pellet was resuspended in hexane and filtered through a 0.45 μm syringe filter to produce the CdSe stock used for shelling.

*CdS shelling:* The CdSe cores were shelled with CdS following protocols previously used by our group and others.^2, 3^ In a 3-neck flask setup on a Schlenk line, 100nmol of the CdSe stock solution, 3 mL OAm, and 3 mL ODE were added. The flask was degassed for ~1 hr with constant stirring (~800 rpm) at room temperature. The contents were then heated to 115 ^o^C and held at this temperature for ~20 min, followed by returning the flask to atmospheric pressure by refilling with N_2_ gas. The contents were then heated to 350 ^o^C at a ramp rate of 16 ^o^C/min, during which time the cadmium and sulfur precursors, 0.150 mmol of Cd-oleate and 0.180 mmol of octanethiol, were prepared in a glovebox. The two precursor solutions were each diluted with ODE to a final volume of 3.5 mL and loaded into two separate syringes. The syringes were then loaded onto a dual syringe pump and injected at a rate of 1.5 mL/hr once the solution reached 200 ^o^C. The solution was held at 350 ^o^C for shell growth, with the final shell thickness determined by taking aliquots every couple of minutes and looking at the shift in the peak position of the PL spectra. Once a desired shell thickness was achieved, the reaction was cooled to 200 ^o^C, followed by dropwise addition of 1 mL OlAc. The solution was allowed to further anneal for 1 hr before cooling to 75 ^o^C, before the contents were transferred to falcon tubes for three rounds of washing with hexane and ethanol. The final product is stored in the glovebox until needed.

#### Aβ(1-16)-PEG-CG synthesis

*Reagents:* Canonical Fmoc- and sidechain-protected amino acids (AAPTec) and Fmoc-8-amino-3,6-dioxaoctanoic acid (Fmoc-PEG, Advanced ChemTech) were purchased and used without further purification. A 100-200 mesh Rink amide resin (cat: SA5013) was purchased from Advanced ChemTech. Activator coupling reagents were purchased from AAPTec (HBTU, cat: CXZ020; HOBt, cat: CXZ010), ApexBio (HOAt, cat: A7024), or Oakwood (HATU, cat: 023926; DIC, cat: M02889). Triisopropylsilane (TIPS, cat: S17975) and trifluoroacetic acid (TFA, cat: 001271) were also purchased from Oakwood. Acetic anhydride (Ac2O, cat: A10), acetonitrile (MeCN, cat: A21), methylchloride (DCM, cat: D35), dimethylformamide (DMF, cat: D119), ethyl ether anhydrous (ether, cat: E138), and methanol (MeOH, cat: A4128) were purchased from Fisher Scientific. N,N-Diisopropylethylamine (DIPEA, cat: D125806) and piperidine (cat: 104094) were purchased from Sigma-Aldrich.

*Peptide Synthesis:* The Aβ(1-16)-PEG-CG sequence, DAEFRHDSGYEVHHQK-PEG-CG, was synthesized following a traditional C-to-N solid phase procedure.^4, 5^ All mixing steps are done on a rotisserie or tube rotator. First 0.1 mmol of Rink resin was loaded into a fritted syringe and swelled with DCM for 5 min on a rotator, followed by filtration and three rounds of rinsing with DMF via vacuum filtration. Deprotection solution (20% v/v piperidine in DMF) is then added to the resin and mixed for 10-15 min, followed by two rinses with DMF. This deprotection step was repeated. The resin is then loaded with the first amino acid G, by two 90 min couplings with HBTU and HOBt in DMF coupling solution containing a 5-molar excess (0.5 mmol) of all components (Fmoc-Gly-OH, HBTU, HOBt). The loaded resin is then capped with an acetic anhydride solution (20% v/v in DMF). Amino acids are then sequentially coupled with HBTU/HOBt as activators in DMF at a 5-molar excess to the rink resin (0.5 mmol) and mixed for an hour. Between each amino acid, the elongating peptide sequence undergoes two rounds of deprotection, comprised of mixing in deprotect solution for 10 minutes followed by two washes with DMF. For the Fmoc-PEG-OH coupling, HATU and HOAt were used as activators in a ~3% v/v DIPEA:DMF solution. Additionally, the lysine (Fmoc-Lys(Boc)-OH) residue coupling following the PEG amino acid was coupled twice to ensure complete strand elongation at all available sites. The completed peptides undergo a final Fmoc deprotection step, followed by cleavage from the resin using a 95:2.5:2.5 v/v/v TFA:TIPS:water solution. The resulting solution was concentrated and the peptide was precipitated via addition of cold ether. The precipitate was collected via centrifugation and dissolved in a 60:40 v/v MeCN:water mixture prior to purification by preparatory HPLC.

*Peptide Purification and Characterization:* Peptide purification was performed using a reverse phase Phenomenex Gemini (5 μm, NX-C18, 110 Å, 250 x 50 mm) column on an Interchim PuriFlash 4125 preparatory HPLC with a binary gradient of water and acetonitrile with 0.1% TFA at 100 ${mL}/{min}$. For fraction collection, UV absorbance of the eluent was monitored at 215 nm and 254 nm. Peptide purity was confirmed using analytical HPLC with a reverse phase Phenomenex Gemini (5 μm, C18, 110 Å, 250 x 4.6 mm) column on a Shimadzu LC-2010A and via MALDI-TOF mass spectroscopy. Peptide concentration for all experiments was determined by analyzing samples via analytical HPLC for comparison to a concentration curve calibrated by amino acid analysis (Figure S2). Appropriate amounts were then aliquoted and lyophilized prior to use.

#### DSPE-PEG_2k_-Aβ conjugation

*Reagents:* All reagents were used as provided by the manufacturers without further purification, unless indicated otherwise in the following protocol. DSPE-PEG_2k_-Maleimide was purchased from BroadPharm (cat: BP-23307). Tris(2-carboxyethyl)phosphine hydrochloride (TCEP) was purchased from Millipore Sigma (cat: C4706). Phosphate buffered saline (PBS) pH 7.2 was purchased from ThermoFisher (cat: 20012027).

*Protocol:* The following details an example reaction scheme involving 1 μmol of the thiol source (Aβ(1-16-PEG-CG) and 1.6 μmol of the maleimide source (DSPE-PEG_2k_-Mal), however the reaction may, to our experience, be scaled up to at least 10x equivalents.

First, any disulfide bridges formed between monomers of Aβ(1-16)-PEG-CG are reduced(?) via incubation of 1 μmol of Aβ(1-16)-PEG-CG with 10-mole excess (10 μmol) TCEP in degassed PBS pH 7.2. Specifically, 2.8 mg of TCEP is added to a 10 mL round bottom flask and 1 μmol of Aβ(1-16)-PEG-CG is dissolved in 8.7 mL of degassed PBS, then added to the flask to dissolve the TCEP. The flask is capped with a septa and punctured with two needles to allow for purging under constant flow of inert gas (N_2_ in this case) while stirring at 400 rpm for 18 hours. An aliquot is taken for analysis via thin layer chromatography ( eluting buffer) to confirm a reaction has a occurred; this is indicated by the presence of a species under short and long UV irradiation with a retention factor different than both the thiol and maleimide sources. After confirmation of a product is formed, the reaction is terminated and the reaction volume is purified via dialysis using a 2k MWCO dialysis cassette (ThermoFisher; cat: ) using a bath solution of nanopure water and three rounds of purification (bath solution was replaced at hours 24, 36, 48?. The purififed product was collected from the dialysis cassette and characterized via analytical HPLC and MALDI-ToF to confirm the observation of a delayed retention peak in HPLC and mass distribution in MALDI-ToF that would be indicative of the product (Figure S2). For MALDI-ToF, the sample was prepared in a matrix composed of α-cyano-4-hydroxycinnamic acid, 0.5% trifluoroacetic acid, and 0.1% NaCl dissolved in EtOH and measured at 60−80% power (Shimadzu Axima Performance). The final DSPE-PEG_2k_-Aβ solution frozen over dry ice, and lyophilized for storage in -20 ^o^C until needed.

#### ABQD Construction

*Protocol:* The DSPE-PEG_2k_-Aβ (0.5 μmol) was dissolved in 0.5 mL of chloroform (CHCl_3_) and combined with 0.05 nmol of CdSe/CdS QDs in a 1 dram vial ( ; cat: ). The mixture was sonicated for 5 minutes at power level 6 (VWR; model 250D) to homogenize the two species and disperse any QD aggregates and inverse micelles of DSPE-PEG_2k_-Aβ. The solution was then dried using a rotary evaporator until a paste like consistency was achieved, to which 500 uL of nanopure water was added to resuspend the solution. Pulse vortexing and sonication (3 minutes at power level 9) were used on occasion to help promote dissolution of the pellet, followed by another round of rotary evaporation to remove any residual organic solvent and drive all QDs into aqueous phase. The ABQD solution should appear transparent. The solution is then stirred for 1 hr at ~400 rpm before undergoing purification via: (i) filtration using a 0.1 μm syringe filter, (ii) ultracentrifugation of the eluent at 100k rpm, 4 ^o^C, 45 minutes, acceleration 5 ( ; model ), (iii) resuspension of the pellet in 100 uL for another round of centrifugation at 16,000 rcf for 30 minutes ( ; model: ), (iv) the supernatant collected and passed through a size exclusion spin column (Cytiva; cat: 27513001) to remove any unencapsulated QDs or free DSPE-PEG_2k_-Aβ. The eluent is the purified ABQD solution that is then characterized via absorbance, photoluminescence, dynamic light scattering (DLS), and transmission electron microscopy (TEM).

*DLS Analysis:* Samples were spotted in duplicate into a 384-well plate ( ; cat: ) and analyzed using a DynaPro system controlled using v7.10 of the DynaPro software (Wyatt). Each well of interest was measured using ten 3 sec acquisitions per run over three runs. The pooled results over analyzed using the “Legacy” settings on DynaPro for fitting. The fitted results were then imported into MATLAB for plotting as the Gaussian-fitted histogram in Figure 1.

*TEM Analysis:* Samples were prepared by spotting 10 uL of the ABQD solution onto an ultrathin lacey carbon-supported copper grids (mesh size 400, Ted Pella; cat: 01824). The solution was allowed to sit for 10 minutes, followed by wicking off using a Whatman #1 filter paper at a 45^o^ angle. This was followed by addition of 10 μL of 1% uranyl acetate as a counterstain. The counterstain was allowed to react for 1 to 2 minutes before wicking off with the same filter paper and drying under a gentle stream of inert gas (N_2_). The grid was then imaged on an FEI TECNAI F-20 field electron microscope with an accelerating voltage of 200 kV.

#### Primary Culture of Hippocampal Neurons and Astroglia

The hippocampal cells were harvested from Sprague-Dawley rats at embryonic day 18. The care and use of these animals abided by the Guide for the Care and Use of Laboratory Animals and followed protocols approved by the University Committee on Animal Resources at the University of Rochester. Hippocampi were dissected and dissociated in 0.25% trypsin (ThermoFisher; cat: 15050065) followed by seeding of the cells at either a density of 45k cells/well on poly-D-lysine coated 12 mm coverslips (Neuvitro; cat: GG-12-15H) in a 24-well plate for general immunocytochemical experiments, or at a density of 120k cells/well on poly-D-lysine coated 25mm coverslips (Structure Probe, Inc.; cat: 01002-AB) for calcium signaling experiments. Cells were first cultured in a supplemented neurobasal media (NBM, ThermoFisher; cat: 21103049) containing 2% B-27^TM^ supplement (ThermoFisher; cat: 17504044), 1% GlutaMAX (ThermoFisher; cat: 35050061), 25 μM glutamic acid, and 5% fetal bovine serum (FBS, Atlas Biologicals; cat: F-0500-D). The culture media was replenished every 3-4 days by aspirating off half of the conditioned media and replacing the aspirated volume with a reduced supplemented NBM containing B-27^TM^ without antioxidants (ThermoFisher; cat: 10889038) and 1% GlutaMAX. Cultures were kept incubated at 37^o^C in a 5% CO_2_ environment.

#### Calcium Transients

*Treatments:* Primary rat hippocampal neuronal glial cultures were treated with 2.5 µM solution of Fluo-4 AM (ThermoFisher; cat: F14201) in NBM for 30 minutes to introduce the cell permeant calcium indicator dye. The Fluo-4 AM solution was made from a stock solution of 2.5 mM Fluo-4 AM in pure DMSO, such that the final 2.5 μM concentration in NBM results in a DMSO content of ≤0.1%. The cells were then washed with NBM for 5 minutes, followed by aspiration and 1 mL treatments of vehicle, 10 nM ABQDs, or 10 nM PEGQDs. These treatments were incubated for 10 minutes, followed by addition of 1 mL KCl in NBM, to result in a total sample volume of 2 mL and final KCl concentration of 30 mM. The KCl addition serves to induce neuronal depolarization and calcium influx.

*Imaging:* Coverslips were transferred to the stage of an inverted Olympus IX-70 microscope equipped with a metal halide arc lamp (X-Cite 120; Excelitas). Samples were imaged 20x magnification (Olympus UPlanApo 20x/0.70 NA objective) on a CCD camera (Q imaging Retiga Exi Fast) using a GFP emission filter (488 TIRF C101581). Images were captured every 20 seconds for 12 minutes with 100 ms exposure times. Cells were placed onto the microscope right after the incubation period of the treatment groups and recordings began as frame 1 being t = 0 sec. After 10 background frames were collected, the 1 mL of KCl in NBM was added. Images were acquired using MetaMorph (Molecular Devices) and saved as .stk files.

*Data Analysis*: All .stk recordings were imported into ImageJ-2 (NIH). Drift and registration was corrected using the default settings on the Linear Stack Alignment with SIFT plugin. The aligned image stacks were then assigned the “Green” lookup table color coding, followed by taking a sub-stack of the first 10 frames. The average intensity of these 10 frames was taken as a representative background signal and subtracted from all frames. The stack was then measured and the recording values were exported as a .csv file for further analysis and plotting on MATLAB. A sample workflow with relevant macro functions are provided as follows:

1. Open file

2. Registration Correction -> Linear Stack Alignment w/ SIFT

run("Linear Stack Alignment with SIFT", "initial_gaussian_blur=1.60 steps_per_scale_octave=3 minimum_image_size=64	maximum_image_size=	1024 feature_descriptor_size=4 feature_descriptor_orientation_bins=8 closest/next_closest_ratio=0.92 maximal_alignment_error=25	inlier_ratio=0.05 expected_transformation=Rigid interpolate");

3. LUT -> Green

run("Green");

4. Substack 1-10

run("Make Substack...", "slices=1-10");

5. Grouped Z Project -> Average Intensity

run("Grouped Z Project...", "projection=[Average Intensity] group=10");

6. Image Calculator -> Subtraction

imageCalculator("Subtract create stack", "______.tif","AVG_Substack (1-10).tif");

7. Measure stack

run("Measure Stack...");

#### BV-2 Microglial Cultures

*Cell Line Maintenance:* BV-2 cells were maintained similarly to as previously described in work from the Gelbard lab relevant to HIV-1 studies of microglial activation.^6^ Specifically, the immortalized murine microglial cells were cultured in Dulbecco’s Modified Eagle Medium (DMEM, high glucose (+) sodium pyruvate (-) L-glutamine; ThermoFisher cat: 10) supplemented with 1% GlutaMAX (ThermoFisher cat: 35050061), 10% fetal bovine serum (FBS; Atlas Biologicals cat: F-0500-D), 1% penicillin and streptomycin (____ cat: ). The cells are plated onto either cell culture flasks, multi-well cell culture dishes, or glass coverslips that are coated with 100 μg/mL poly-D-lysine (PDL) in ddH_2_O via room temperature incubation for 1-3 hours, followed by three washes with ddH_2_O. For glass coverslips, 20 minutes of plasma cleaning is additionally done prior to PDL deposition to ensure sterility of the surface. The cultures are passaged between 80-90% confluency (roughly 2-3 days) by digestion of surface adhesion proteins via 0.25% trypsin-EDTA (ThermoFisher cat: 25200056) for ~5 minutes in a 37 ^o^C + 5% CO_2_ cell culture incubator. Cell were passaged into fresh T-175 culture flasks at a density of ~3 million cells/flask, 24-well cell culture plates (with and without coverslips) at 50k cells/well, and 12-well cell cultures plates at 85k-100k cells/well.

*Treatment of BV-2 cells:* Prior to treatment of the BV-2 cultures at ~75% confluency, the cells were first serum deprived for 45 minutes in the same complete DMEM composition as for culturing, except at a reduced serum content of 1% FBS to prime the cells to be in a similar cell cycle to minimize heterogeneity within a single well. The reduced serum media was then gently aspirated off and replaced with the following treatment groups dissolved in the same reduced serum DMEM: vehicle (0.1% DMSO), 100 nM URMC-099 (from 100 μM stock in 100% DMSO), 50 nM ABQDs, 50 nM PEGQDs, 100 nM URMC-099 + 50 nM PEGQDs, 100 nM URMC-099 + 50 nM ABQDs. After ~18 hours, the treatments were aspirated off, briefly rinsed with ice cold Dulbecco’s phosphate buffered saline (DPBS; ThermoFisher cat: ) and processed either for RT-qPCR or immunofluorescent imaging following the protocols described in the following sections.

#### RT-qPCR

*RNA isolation:* RNA as isolated from BV-2 microglia using a PureLink^TM^ RNA Mini Kit (Invitrogen cat: 12183018A). After isolation, the purity (A260/A280 and A260/A230) and concentration (A260) of the extracted RNA was determined using a NanoDrop^TM^ Spectrophotometer (Thermo Scientific). The total RNA was then stored at -80 ^o^C until cDNA synthesis for downstream applications.

*cDNA synthesis:* Total RNA from the BV-2 microglia was used to generate cDNA using the SuperScript III First Strand Synthesis Kit (Invitrogen cat: 18080051), using 1 μg of total RNA as a template. The final cDNA was measured for purity and concentration on the same NanoDrop^TM^ Spectrophotometer and stored at -20 ^o^C until RT-qPCR analysis.

*RT-qPCR measurement:* RT-qPCR was performed using specific primers designed to target inflammation (CXCL10) and unfolded protein response (CHOP, XBP-1 splicing) pathways (Table S1). The SYBR GreenER^TM^ qPCR SuperMix Universal (Invitrogen cat: 11762100).

#### Immunofluorescent Staining

The neuroglial co-cultures and BV-2 microglia cultures on coverslips were stained using an indirect immunofluorescent labeling protocol. The pairings of primary and secondary antibodies are outlined in Tables S2 and S3. After staining, the coverslips were mounted onto cover glass and stored protected from light until needed for imaging.

#### OptiGrid Structured Illumination Imaging and Analysis

*Fluorescent Imaging w/ “Grid” Confocal Microscope:* The immunostained samples were imaged on an Olympus BX51 microscope excitation with a Prior Lumen 200 illumination source equipped with a Hg lamp (Prior cat: LM200B1-A) and detected with a Hamatsu ORCA-ER scientific camera. The following pairs of excitation and emission filters were used to selectively illuminate and detect biomolecular targets immunolabeled with DAPI, AlexaFluor488, AlexaFluor568, AlexaFluor647, and CdSe/CdS QD, respectively: 350 nm / DAPI (Semrock; cat: FF02-447/60-25); 405 nm / FITC (Semrock; cat: FF01-524/24-25); 488 nm / TRITC (Semrock; cat: FF01-593/40-25); 568 nm / Cy5 (Semrock; cat: FF01-692/40-25); 350 nm / TRITC. An infinity-corrected 20x UPlanApo 0.70 NA objective (Olympus) was used to collect and collimate the emission through an OptiGrid structured illumination element to form a “grid” confocal image on the detector. Z-stacks were captured for all samples at 1 μm step sizes and compressed into the extended focus view presented in the representative images used in this manuscript. The exposure time for each channel was optimized and kept the same between each sample.

*Volocity Image Analysis:* Volocity 3D Image Analysis software (PerkinElmer) was used to analyze the collected image sets. The general workflow of A fine noise filter was used on all images before applying a set of measurement protocols. For the neuroglial cultures, a measurement protocol was designed to identify objects above a certain threshold corresponding to immunofluorescent labeled nuclei (λ_ex_ = 350 nm, DAPI BPF), PSD-95 (λ_ex_ = 405 nm, FITC BPF), MAP2 (λ_ex_ = 488 nm, TRITC BPF), and GFAP (λ_ex_ = 568 nm, Cy5 BPF), as well as CdSe/CdS QD emission (λ_ex_ = 350 nm, TRITC BPF). The sum of measured object intensities and spatial volume was extracted for the PSD-95, MAP-2, and GFAP objects. The MAP-2 objects were also further analyzed to extract the prevalence of dendritic beading by setting a cutoff for object volume and threshold for spheroidicity to identify true “beads.” The total number of beads, nuclei, and CdSe/CdS objects were also exported for further analysis.

#### Statistical Analysis

All quantitative values were organized and pre-processed on Excel prior to importing the values onto GraphPad Prism 9. Each replicate value was imported. For the rescue experiments of the NVU co-culture experiments, a two-way ANOVA with Holm-Sidak post-hoc correction was used. For all other experiments, a one-way ANOVA with Holm-Sidak post-hoc correction was used. Statistical significance was defined as an adjusted p-value less than 0.05 for all analyses.

### Schemes

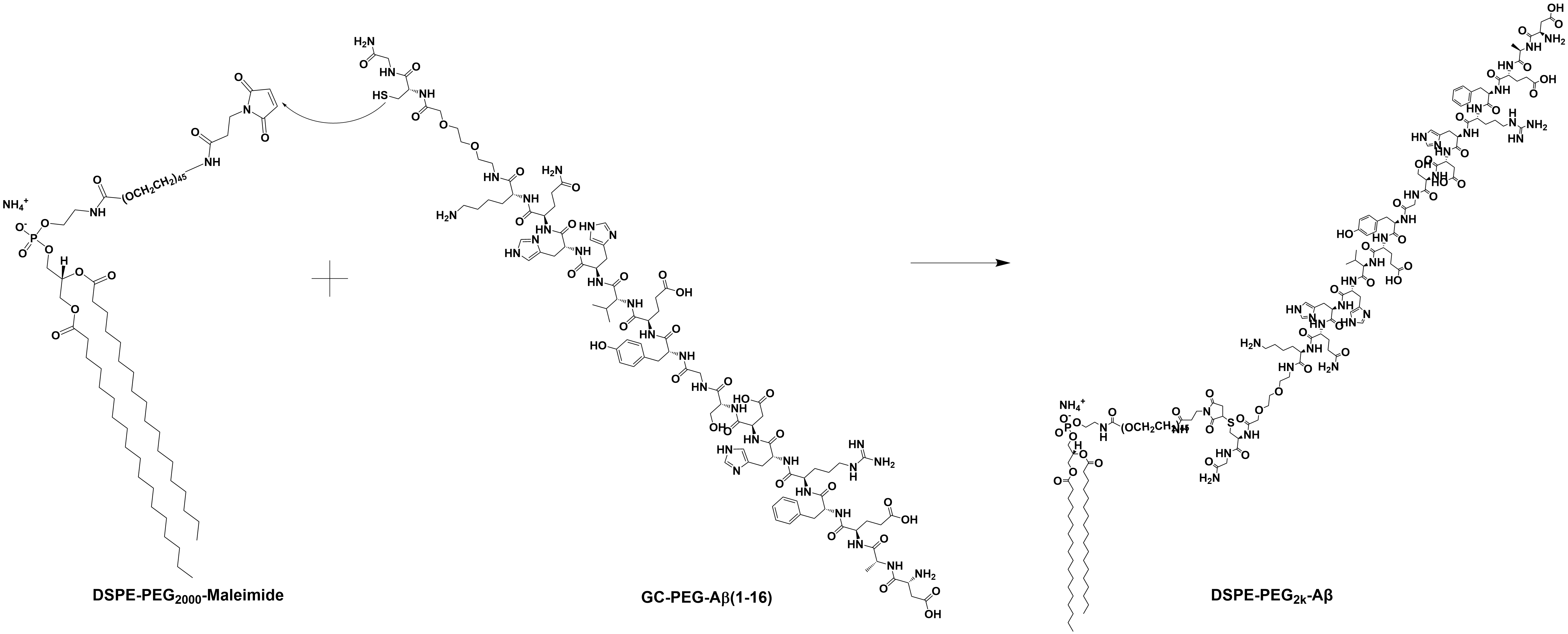

Scheme S1. Thiol-maleimide conjugation reaction between DSPE-PEG_2k_-Mal and Aβ(1-16)-PEG-CG to produce DSPE-PEG_2k_-Aβ.

Sulfhydryl of cysteine in Aβ(1-16)-PEG-CG acts as a nucleophilic donor towards the maleimide (olefin) ring in DSPE-PEG_2k_-Maleimide to allow for a Michael addition mediated “click” chemistry to form the peptide-functionalized phospholipid PEG.

### FIGURES

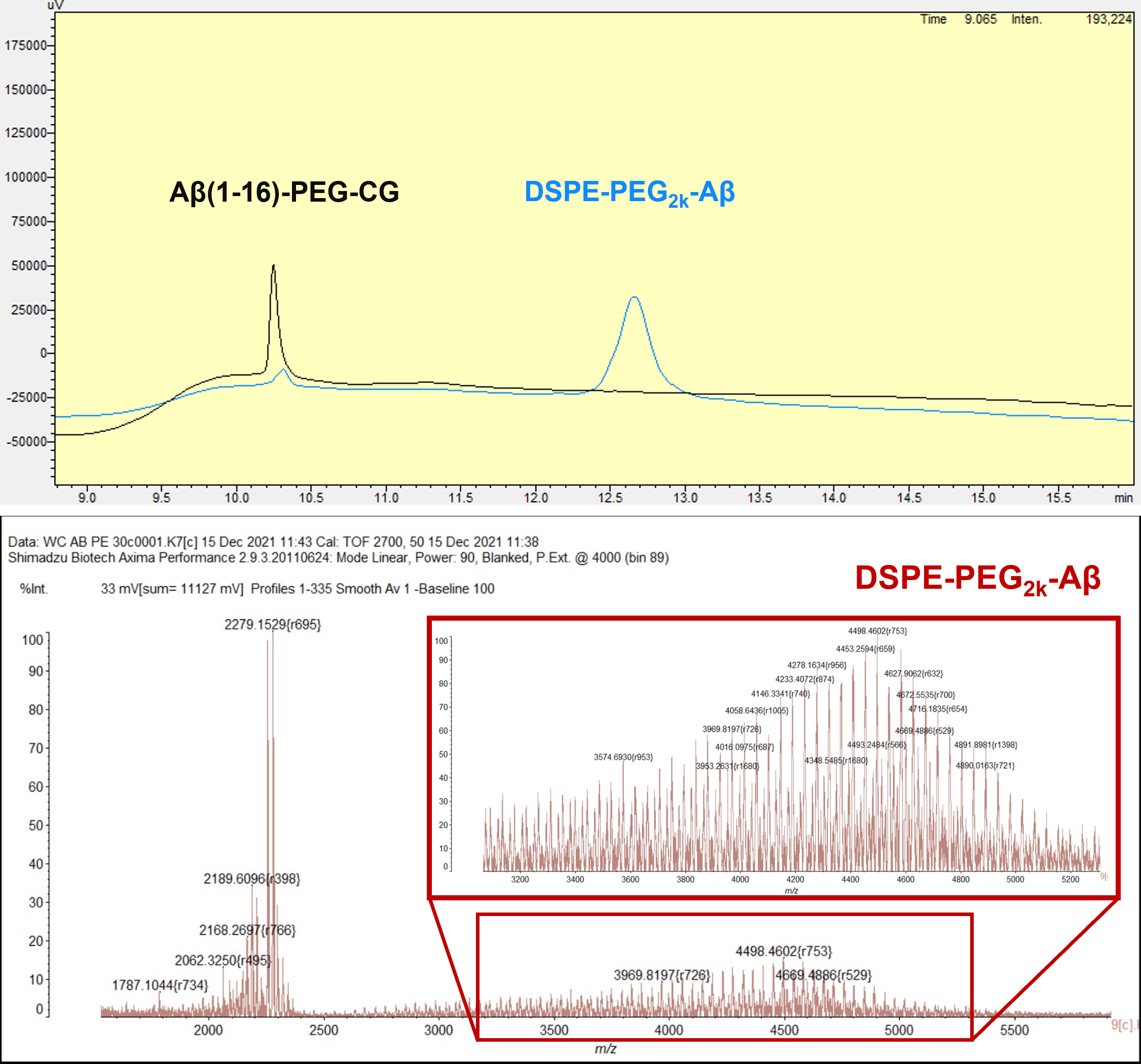

Figure S1. HPLC (top) and MALDI-ToF (bottom) spectra to confirm formation of DSPE-PEG_2k_-Aβ.

This is evidenced by the formation of a delayed retention peak in the HPLC spectra (second peak) in the DSPE-PEG_2k_-Aβ spectra (blue) compared to the pure Aβ(1-16)-PEG-CG curve (black). This is complemented by the same sample being spotted for MALDI-ToF, resulting in the observation of a broad lipid PEG distribution centered at ~4300, reflective of DSPE-PEG_2k_-Aβ formation due to the combination of the Aβ(1-16)-PEG-CG (sharp peaks at 2279) and DSPE-PEG_2k_-Maleimide (broad distribution centered at 2000).

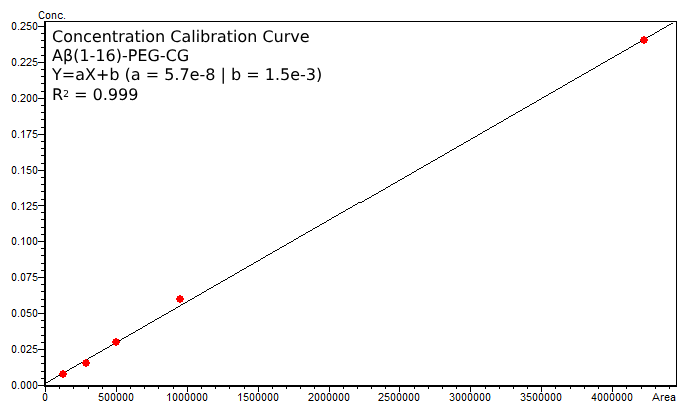

Figure S2. Analytical HPLC calibration curve for concentration determination of Aβ(1-16)-PEG-CG.

Figure S3. PEGQD treated neuroglia

Figure S4. PEGQD phagocytosis by BV-2

Figure S5. CdSe/CdS QD spectra to ABQD spectra (Abs Norm PL)

### TABLES

| Table S1. Selection of primary antibodies and dilutions used for immunofluorescent staining | | | | | |
| --- | --- | --- | --- | --- | --- |
| Culture | Host | Target | Dilution | Source | Catalog # |
| BV-2 | Rabbit | Iba1 | 1:500 | Wako | 019-19741 |
|  | Rat | Lamp1 | 1:500 | ThermoFisher | 14-1071-82 |
| neurons | Mouse | PSD95 | 1:200 | NeuroMab | 75-028 |
|  | Rabbit | MAP2 | 1:150 | Cell Signaling | 4542 |
| astrocytes | Chicken | GFAP | 1:250 | Neuromics | CH22102 |

| Table S2. Selection of secondary antibodies used for immunofluorescent staining | | | | | |
| --- | --- | --- | --- | --- | --- |
| Host | Reactivity | Fluor | Dilution | Source | Catalog |
| Goat | Rabbit | Alexa 568 | 1:750 | ThermoFisher | A-11036 |
|  | Rat | Alexa 647 |  | ThermoFisher | A-21247 |
|  | Mouse | Alex 488 |  | ThermoFisher | A-11006 |
|  | Chicken | Alexa 647 |  | ThermoFisher | A-21449 |
